# Genome-Wide Identification of Essential Genes in the Invasive *Streptococcus anginosus* Strain

**DOI:** 10.1101/2025.06.09.658608

**Authors:** Aleksandra Kuryłek, Jan Gawor, Karolina Żuchniewicz, Robert Gromadka, Izabela Kern-Zdanowicz

## Abstract

**Background:** *Streptococcus anginosus*, part of the *Streptococcus anginosus* group (SAG), is a human commensal increasingly recognized as an opportunistic pathogen responsible for abscesses formation and infections, also invasive ones. Despite its growing clinical importance, the genetic determinants of its pathogenicity remain poorly understood.

**Objectives:** This study aimed to identify essential genes in *S. anginosus* 980/01, a bloodstream isolate, under optimal laboratory conditions using a transposon mutagenesis combined with Transposon-Directed Insertion Site Sequencing (TraDIS).

**Methods:** A mutant library was generated using the IS*S1* transposon delivered *via* the thermosensitive plasmid pGh9:IS*S1*. Following transposition, insertions were mapped using Illumina sequencing and analyzed. Essential genes were identified based on the absence of insertions and statistical filtering.

**Results:** The library exhibited 98% genome saturation with over 130,000 unique insertion sites. Among 1,825 genes, 348 (19.1%) were essential, 1,446 non-essential, and 30 non-conclusive. Comparative analyses were performed with *S. pyogenes* MGAS5005 and *S. agalactiae* A909. Similarly to the latter, essential genes were enriched in functions related to translation, transcription, and cell wall biosynthesis. However, 40 genes uniquely essential to *S. anginosus* 980/01 were identified, suggesting unique survival strategies in *S. anginosus*.

**Conclusions:** This study presents the first genome-wide identification of essential genes for *S. anginosus* 980/01, highlighting conserved and unique essential genes. These findings provide a basis for understanding its pathogenic potential and for identifying novel antimicrobial targets.

## Introduction

*Streptococcus anginosus,* together with *S. constellatus* and *S. intermedius,* constitute the *Streptococcus anginosus* group (SAG) commonly found on mucosal membranes in the healthy human microbiota, particularly in the oral cavity, gastrointestinal tract, and urogenital tracts [1]. Despite this, SAG species are increasingly recognized as opportunistic pathogens, especially in immunocompromised or cystic fibrosis patients [2–4]. Among SAGs, *S. anginosus* is frequently isolated from abscesses, bloodstream infections, and polymicrobial infections, and is more commonly associated with invasive infections than other SAG members, also in individuals without any underlying diseases [2,5–8].

Interestingly, *S. anginosus* strains exhibit variable hemolytic activity and can express different Lancefield antigens, complicating diagnosis and classification. These strains can also produce diverse virulence factors [9,10]. However, the molecular mechanisms underlying the *S. anginosus* pathogenesis remain poorly understood, with many of its genes still annotated as hypothetical [7,11,12]. To assess the essentiality and fitness of each gene within the bacterial genome, the most efficient strategies integrate the next-generation sequencing (NGS) technologies with large-scale transposon mutagenesis [13–17]. This approach enables the identification of the chromosomal sequences adjacent to transposon integration sites. However, genes required for bacterial growth and survival under specific experimental conditions, termed essential genes, cannot be disrupted. A saturated transposon mutant library contains a collection of viable mutants with a transposon inserted into every non-essential gene, while insertions into essential genes are lethal. Moreover, such a library can be challenged in specific conditions, such as human blood or antibiotics, to distinguish non-essential genes that are necessary for bacterial survival and growth in this particular environment.

For mutagenesis of Gram-positive bacteria, transposons are typically based either on *mariner* [18–21] or IS*S1* [22–26], both utilizing a copy-out–paste-in mechanism. While the *mariner*-family transposons preferentially integrate at TA dinucleotide sequences [27], the IS*S1* does not exhibit such specificity [22]. Essential genes of several streptococcal species have been analysed in the IS*S1* Transposon-Directed Insertion Site Sequencing (TraDIS), including *S. uberis* 0140J [17], *S. equi* 4047 [24], and *S. agalactiae* CNCTC 10/84 [28]. Meanwhile, for *S. pyogenes* M1T1 5448 [18], *S. agalactiae* A909 [29], and *S. suis* SC19 [30], the *mariner*-based Krmit or Himar1 have been used. The identification of essential genes provides promising targets for the development of novel antimicrobial therapies [31].

In this study, we performed an IS*S1* TraDIS approach to identify genes essential for the growth of *S. anginosus* 980/01 [11], an invasive strain isolated from sepsis, under optimal laboratory conditions. As the first genome-wide transposon mutagenesis study in *S. anginosus*, this analysis provides an initial framework for exploring its essential gene set. The TraDIS data for *S. anginosus* were finally compared to those of *S. pyogenes* and *S. agalactiae* to identify conserved essential genes and determine the unique set of genes essential to *S. anginosus* 980/01. The transposon-based methods, such as TraDIS, have become powerful tools for defining the genetic determinants of bacterial survival and virulence.

## Materials and Methods

The *S. anginosus* 980/01 strain, isolated from the bloodstream of a 67-year-old patient in 2001 in Poland, exhibits α-hemolytic activity and expresses the Lancefield group F antigen on its surface. The strain is part of the Collection of the National Institute of Medicines, Warsaw, Poland [11].

### S. anginosus cultures

The *S. anginosus* was grown on BHI agar (VWR chemicals) plates or in BHI broth (VWR chemicals) supplemented, if necessary, with erythromycin (Acros Organics) in a concentration of 5 µg/ml. Bacteria were grown at 37°C, unless otherwise stated, in the presence of a CO_2_ generator, CO_2_Gen (Thermo Fisher Scientific, Waltham, USA). For extended storage, *S. anginosus* 980/01 was frozen in BHI with 15% glycerol at -80°C.

### *S. anginosus* electrocompetent cells preparation

*S. anginosus* 980/01 was grown on BHI agar for 24 hours. After that, the 25-fold diluted culture was grown in BHI to OD₆₆₀ ∼0.3, harvested by centrifugation (5000 × g, 4°C), washed three times with ice-cold 0.5 M sucrose, and resuspended in 0.5 M sucrose with 15% glycerol. Aliquots (50 μL) were stored at –80°C.

### Transformation of *S. anginosus* with the pGh9:IS*S1* plasmid

The pGh9:IS*S1* plasmid [22] with thermosensitive replication was used as a donor of the IS*S1* transposon. Electrocompetent *S. anginosus* 980/01 cells were transformed with 50 ng of pGh9:IS*S1* DNA via electroporation (2.5 kV/cm, 200 Ω, 25 μF, 5 ms pulse; Bio-Rad Gene Pulser). Cells were recovered in BHI at 28 °C for 4 h; after 1 h of incubation, erythromycin in a sublethal concentration was added to induce resistance expression. Transformants were selected on BHI with erythromycin (BHIE) at 28°C after 48 h-incubation.

### The *S. anginosus* 980/01 mutant library construction

A single colony of *S. anginosus* 980/01 carrying pGh9:IS*S1* was grown in 10 mL BHIE at 28°C for 24 h, then heat-shocked at 40°C for 2.5 h to stop plasmid replication and induce IS*S1* transposition. Transposants were selected after overnight growth on 100 BHIE plates (∼6,500 colonies/plate) at 40°C, in an atmosphere with 5% CO₂. Colonies were harvested into BHI with 15% glycerol and stored at −80°C.

### PCR detection of pGh9:IS*S1*

To confirm the presence of pGh9:IS*S1* in selected clones, PCR was performed using DreamTaq DNA polymerase and dNTPs (Thermo Fisher Scientific), with FwIS*S1* and RvIS*S1* primers **(Supplemental File S1**). The reaction conditions: initial denaturation at 95°C for 3 min; 30 cycles of 95°C for 30 s (denaturation), 50°C for 30 s (annealing), and 72°C for 4 min (extension); followed by a final extension at 72°C for 7 min. PCR products were analyzed by agarose gel electrophoresis.

### Identification of IS*S1* insertion sites in mutants

To verify the IS*S1* transposon insertion site in an individual mutant, genomic DNA was extracted and digested with FastDigest HindIII (Thermo Fisher Scientific). The resulting fragments were ligated using T4 DNA Ligase (Thermo Fisher Scientific), and served as templates for PCR amplification with DreamTaq DNA Polymerase (Thermo Fisher Scientific) using primers FwSAGIS*S1* and RvSAGIS*S1* (**Supplemental File S1**). The PCR products were separated by agarose gel electrophoresis, the bands were excised, purified (MicroElute Gel Extraction Kit, OMEGA Bio-tek, USA), and Sanger sequenced using FwSAGIS*S1* (detailed results are provided in **Supplemental File S1**).

### IS*S1* transposon mutant libraries prepration for Illumina sequencing

The portion of IS*S1* transposon mutant library was regrown to OD_660_ = 0.3 in BHIE, spread onto 25 BHIE plates each, pooled, and harvested prior DNA extraction. Genomic DNA was isolated from the bacterial mutant pools using the SDS/Phenol method as described previously [32,33]. DNA quality control was performed by measuring the absorbance at 260/230, template concentration was determined using the Qubit fluorimeter (Thermo Fisher Scientific). DNA integrity was analyzed by 0.8% agarose gel electrophoresis.

The IS*S1* transposon mutant libraries were constructed according to the protocol [Transposon insertion sequencing (Tn-seq) library preparation protocol – includes Unique Molecular Identifiers (UMI) for PCR duplicate removal: https://www.protocols.io/view/transposon-insertion-sequencing-tn-seq-library-pre-rm7vzn6d5vx1/v1] with minor modifications. In brief, DNA was mechanically sheared using Covaris (Covaris, MA, USA) into 300-500 bp fragment sizes, followed by end repair and TA-adaptor ligation using NEB Ultra II End Repair and Ligation Modules (New England Biolabs, Beverly, USA). Ligation reaction was purified by Ampure XP magnetic beads (Beckman Coulter, Brea, USA) and library restriction digest was perfomed using SmaI (Thermo Fisher Scientific) for 2 h at 25°C to cleave the pGH9:IS*S1* plasmid 33 bp upstream of the sequence encoding IS*S1* to minimise the amount of TnSeq reads mapping to plasmid. The digested library was purified using Ampure XP beads, and the amount of DNA recovered was quantified using the Qubit dsDNA HS assay kit (Thermo Fisher Scientific) according to the manufacturer’s instructions. One hundred nanograms of library DNA was PCR amplified for 20 cycles according to the NEBNext Ultra II DNA library prep kit protocol. Library amplification utilized the specific IS*S1* primer containing Nextera XT 5’ overhang (P5) with barcode index sequence and a TA–adaptor compatible unique indexing Nextera XT PCR (P7) primer per library, which facilitated the attachment of the final product to the sequencing flow cell (primer sequences are provided in **Supplemental File S1**). The libraries were quality-checked using the KAPA Library Quantification kit (KAPA-Roche, Basel, Switzerland), pooled in equimolar ratio, and sequenced on a NextSeq 550 instrument using the NextSeq HighOutput reagent v2.5 (150 cycle) chemistry kit (Illumina, San Diego, USA).

### TnSeq data analysis

Raw demultiplexed fastq files were analysed using the TnSeq UMI scripts (https://github.com/nppalani/TnSeq/ and https://github.com/apredeus/TRADIS) modified to handle IS*S1* transposon sequence containing data. Initially, the single command pipeline script, fastqtoreadcount_umi.sh, was run. The pipeline performed UMI-based PCR duplication removal, filtered and removed reads according to the transposon tag. After tag removal, the remaining reads were mapped to the *S. anginosus* 980/01 reference genome. Finally, raw read counts per insertion position output files were generated. Transposon insertions were viewed in Integrative Genomics Viewer [34]. TnSeq data analysis was further performed using Transit v.3.2.3 [35]. Gene essentiality was calculated using the Tn5Gaps method (https://transit.readthedocs.io/en/latest/method_tn5gaps.html), designed for transposons with random genome insertion. This method identifies the longest regions in the gene without insertions to estimate gene significance. The IS*S1* insertion index was calculated for each gene ([Number of unique insertion sites in gene] / [Gene length in bp]). Based on it, the genes were considered essential, non-essential, and non-conclusive. Additionally, for genes considered essential, a statistical significance threshold of p<0.05 was used after applying the FDR (false discovery rate method) to minimize the occurrence of false positives.

## Results

### Genome structure and mobile genetic elements of *S. anginosus* 980/01

The *S. anginosus* 980/01 genomic DNA was previously sequenced using both short-reads of Illumina and the long-reads of Oxford Nanopore technology to obtain the complete physical map; the sequencing of this strain is a part of the larger project and will be described elsewhere. It consists of 1,883,767 bps chromosome, with no plasmids. The sequence is deposited in GenBank under accession number: CP183189, and as BioProject No.PRJNA1228600. The mean %G+C content is 39%. Genome annotation using DFAST v.1.2.18 software (https://github.com/nigyta/dfast_core) revealed that the genome contains 1825 genes coding for proteins, of which 415 are of unknown function. No antimicrobial resistance genes were detected using ResFinder v.4.3.3 (https://orbit.dtu.dk/en/publications/resfinder-an-open-online-resource-for-identification-of-antimicro). Nine putative streptococcal virulence factors were identified using ABRicate v.1.0.1 (https://github.com/tseemann/abricate) with the Virulence Factor Database (https://www.mgc.ac.cn/VFs/), retaining only hits with >80% both sequence identity and target coverage (**Supplemental File S1**). Detection of mobile genetic elements revealed: a/ an intact single prophage, 38.9 kb in size (position 591,476 – 630,426), detected using Phastest (https://phastest.ca/), b/ two putative integrative and conjugative elements (ICE) as well as c/ a single putative integrative and mobilizable element (IME); ICEs and IME were identified using both ICEfinder (https://bioinfo-mml.sjtu.edu.cn/ICEfinder/ICEfinder.html) and ICEscreen v1.3.3 (https://icescreen.migale.inrae.fr/) (**Supplemental File S1**).

### The *S. anginosus* 980/01 mutant library construction

To construct the *S. anginosus* 980/01 transposon mutant library, the pGh9:IS*S1* plasmid of thermosensitive replication origin was used as a donor of the IS*S1* transposon. To enable IS*S1* transposition, the bacterial culture was incubated at a restrictive temperature as described in Materials and Methods section. Due to replicative IS*S1* transposition into the random chromosomal site, the plasmid was incorporated between two duplicated IS*S1* transposon sequences (**Fig. 1**). The resulting library contained 5.3 × 10^9^ clones, with the IS*S1* transposition frequency of 0.29% (calculated as CFU on BHIE versus BHI).

**Figure 1.**
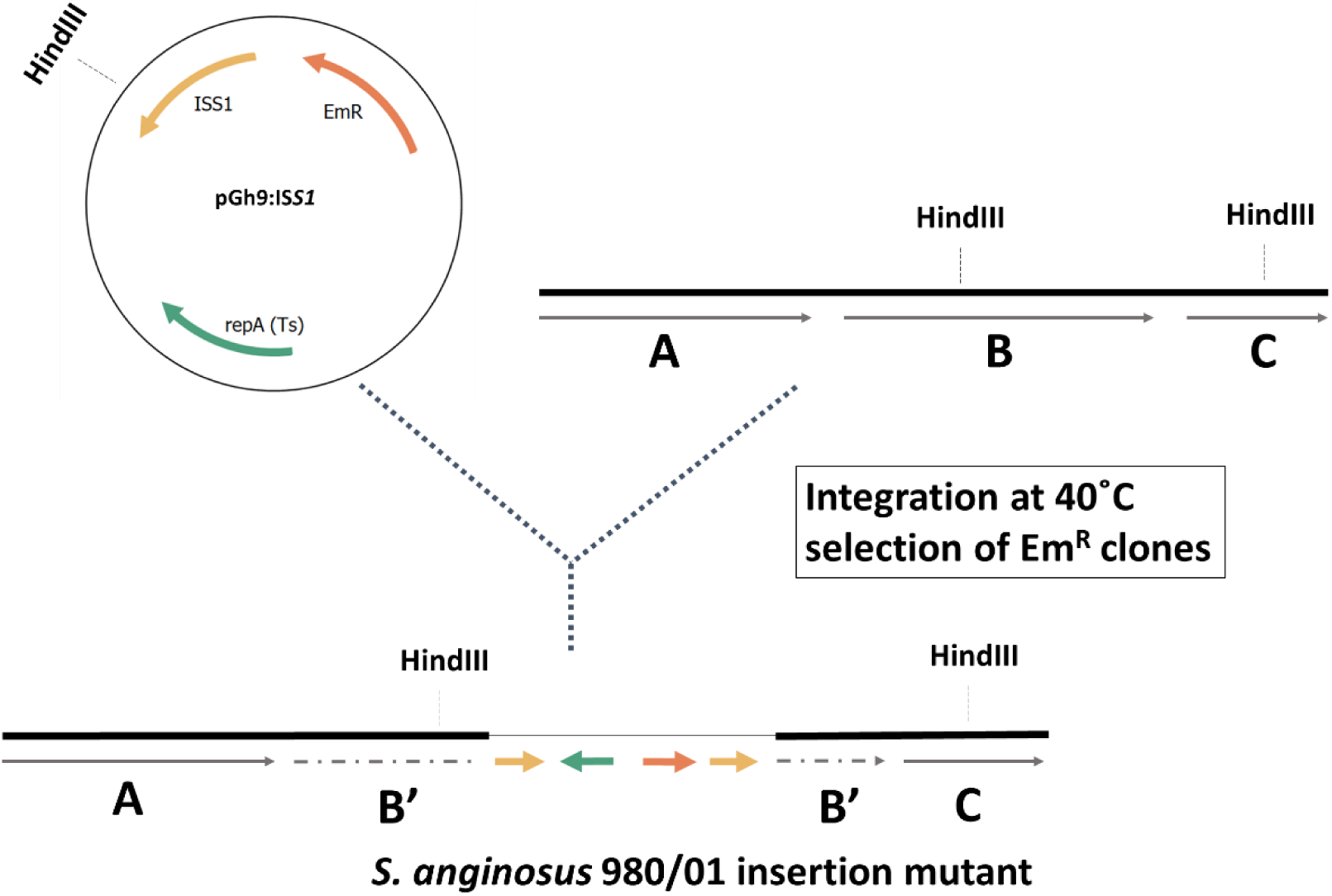
Schematic representation of IS*S1* transposition into the *S. anginosus* 980/01 chromosome. The *S.anginosus* chromosome is shown as a thick black line, while pGh9:IS*S1* as a thin black line. Arrows indicate gene localisation.

### The *S. anginosus* 980/01 mutant library validation

The IS*S1* integration sites in 30 randomly chosen erythromycin-resistant clones from the IS*S1 S. anginosus* 980/01 mutant library were analysed. The sequencing results revealed that in 28 out of 30 clones, IS*S1* was integrated into unique sites, whereas in two mutants (6.7%), IS*S1* was detected at redundant genomic locations. Nevertheless, the library predominantly comprises unique clones, with 93.3% showing distinct integration sites. (**Table 1**).

**Table 1:**
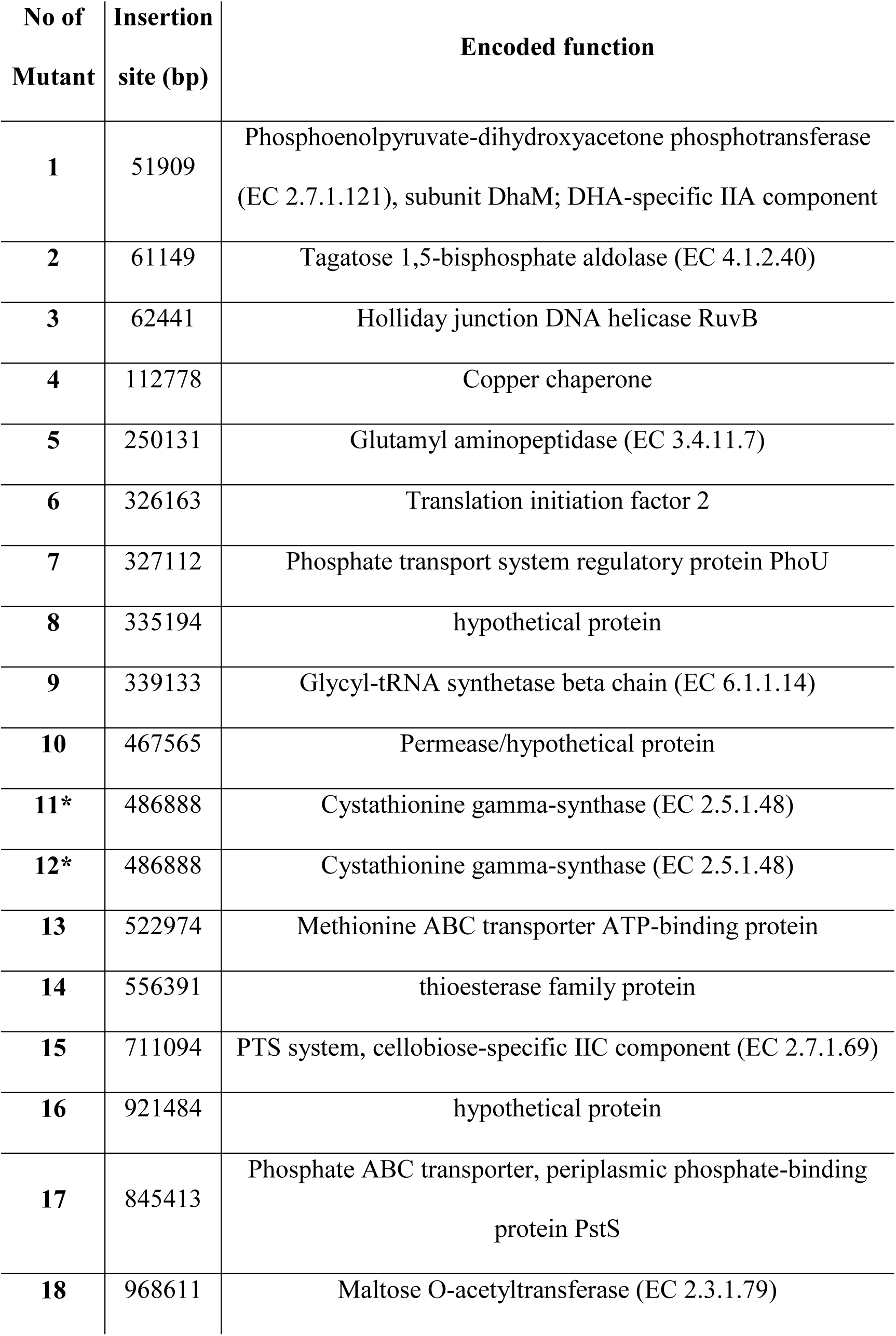
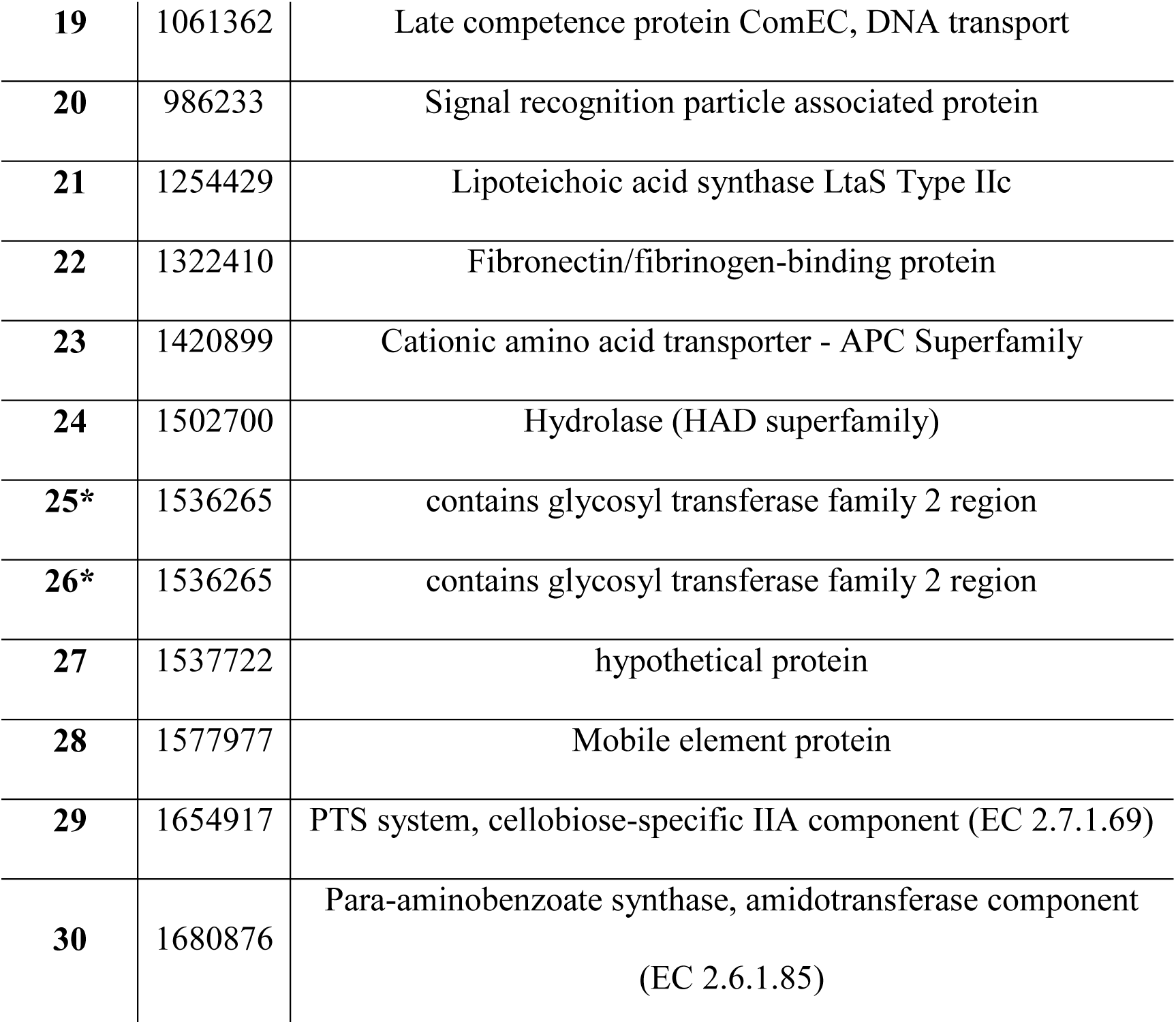
Transposon insertion sites in 30 mutants. The numbers of the redundant mutants are marked with an asterisk (*).

No statistically significant differences in the growth rates were observed between *S. anginosus* 980/01 WT, *S. anginosus* 980/01 carrying pGh9:IS*S1*, and the *S. anginosus* 980/01 transposon mutant library. (**Supplemental File S2**).

### The *S. anginosus* 980/01 mutant library characteristics

For sequencing, three independent replicates of the IS*S1* transposon mutant library (BHI_A, BHI_B, and BHI_C) were regrown and prepared for DNA extraction. After sequencing, each of them was analyzed using the Transit software with the Tn5gaps method [35].

Analysis revealed 132,000 to 175,000 single insertion sites of IS*S1* in the genome, on average every 10.7-14.3 nt. This corresponds to a library saturation of 98% (1,796 disrupted genes of 1,825 annotated), with an average of 64.7 insertions per disrupted gene and a mean of 2,718 sequencing reads per gene (**Supplemental File S1**). The IS*S1* insertion index calculated for each gene was a base to consider a given gene essential, non-essential, and non-conclusive (**Supplemental File S1**). For *S. anginosus* 980/01 essential genes, the FDR method was applied.

Based on these parameters, 348 of 1825 (19.1%) of the *S. anginosus* 980/01 genes were classified as essential, 1446 (79.2%) were non-essential, while 30 (1.7%) were non-conclusive. The functional categories analysis of essential genes is presented in **Fig. 2**. The main category encompasses genes involved in translation, transcription, and cell wall biogenesis.

**Figure 2.**
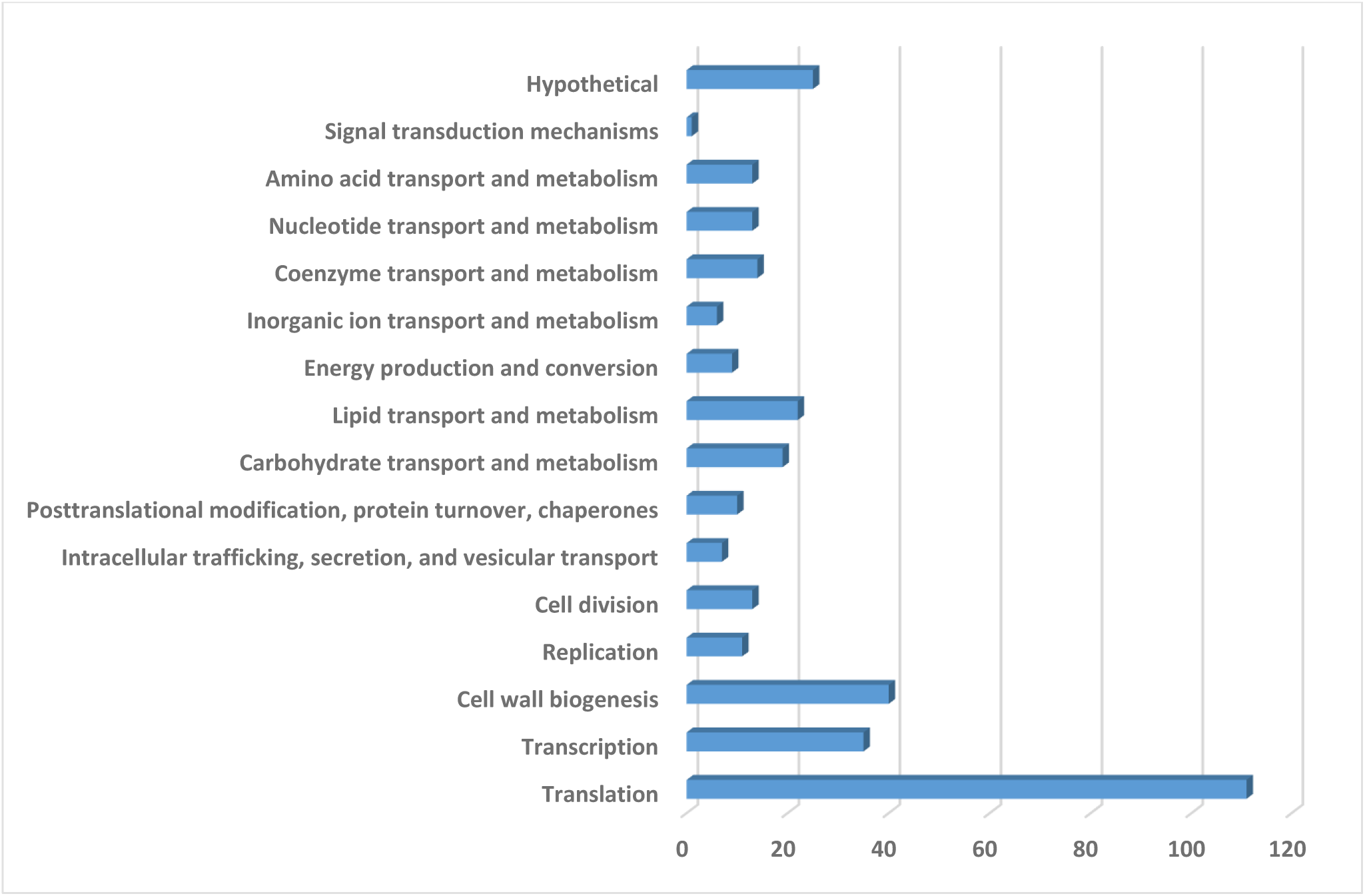
The functional classification of 348 genes essential for *S. anginosus* 980/01.

### Reconstruction of metabolic pathways

Based on the whole-genome analysis of *S. anginosus* 980/01, metabolic pathways were reconstructed using the KEGG Mapper of Kyoto Encyclopedia of Genes and Genomes (https://www.kegg.jp). They were compared with data for *S. pyogenes* M1T1 5448 [36] and *S. agalactiae* A909 [37], other human pathogenic streptococci. Like *S. anginosus* 980/01, these strains were originally isolated from the blood of patients with sepsis. Notably, the study on *S. pyogenes* M1T1 5448 relied on the genome annotation of *S. pyogenes* MGAS5005, as the genome sequence of M1T1 5448 had not yet been published.

The comparison of KEGG data indicates that elements involved in DNA replication and repair, transcription, and translation, as well as the major metabolic pathways, are similar to those of *S. pyogenes* MGAS5005 and *S. agalactiae* A909. So, for ATP production, *S. anginosus* 980/01 relies on fermentation and substrate phosphorylation.

In predicted carbohydrate metabolism, due to the presence of specific genes, *S. anginosus* 980/01 can carry out the citrate cycle reactions facilitated by citrate synthase, aconitate hydratase, and isocitrate dehydrogenase. Those genes are absent in *S. agalactiae* A909 and *S. pyogenes* MGAS5005, but present in e.g., *S. suis* and *S. mutans* [38]. However, these genes are not essential for S*. anginosus* 980/01.

### Comparison of essential gene sets of *S. anginosus* 980/01 with *S. pyogenes* and *S. agalactiae*

To identify conserved and species-specific essential genes, we compared the *S. anginosus* 980/01 dataset with published essential gene sets from other human pathogenic streptococci, *S. pyogenes* M1T1 5448 [36] and *S. agalactiae* A909 [37]. Like *S. anginosus* 980/01, these strains were originally isolated from the blood of patients with sepsis.

For this analysis, the data from Le Breton et al. and Hooven et al. [18,29] were utilized. Genes from *S. pyogenes* and *S. agalactiae* were categorized by the authors as essential, critical, non-essential, or not-defined/inconclusive. Orthologs were identified using OrthoFinder (https://github.com/davidemms/OrthoFinder) and comparisons were made between: a/*S. anginosus* 980/01 and *S. pyogenes* MGAS5005, b/ *S. anginosus* 980/01 and *S. agalactiae* A909, and c/ *S. pyogenes* MGAS5005 and *S. agalactiae* A909.

The essentiality classifications of each orthologous pair were compared, excluding genes with the essentiality unknown or inconclusive. The results are summarized in **Fig. 3** and **Supplemental File S1**.

**Figure 3.**
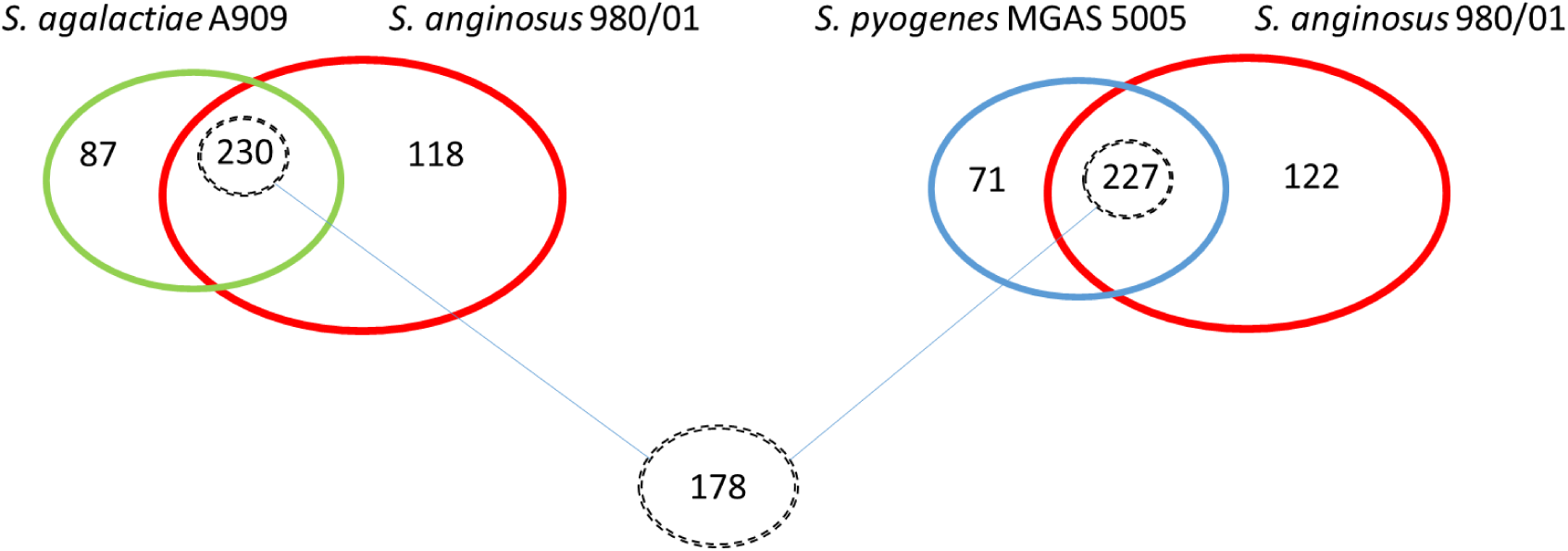
The number of essential genes shared between *S. anginosus* 980/01, *S. pyogenes* MGAS5005 and *S. agalactiae* A909.

The proportion of essential genes in *S. anginosus* (19%) is comparable to that in *S. pyogenes* MGAS5005 (298 essential genes of 1866; 16%) [18] and *S. agalactiae* A909 (317 of 2136; 15%) [29]. Among the genes classified as essential in all three species, 178 (53.5%) were shared; between *S. anginosus* and *S. pyogenes* it was 59.9%, and 62.4% between *S. anginosus* and *S. agalactiae*. These 178 genes essential for 3 strains were enriched in categories such as translation (53 genes), cell wall biogenesis (19), lipid transport and metabolism (16), replication and amino acid transport and metabolism (13 each), carbohydrate transport and metabolism (12), coenzyme transport and metabolism (9), transcription (8), and energy production and conversion (6). Other functional groups included posttranslational modification, protein turnover, and chaperones, inorganic ion transport and metabolism, intracellular trafficking and nucleotide transport and metabolism (5 genes each), cell division, secretion, and vesicular transport (4 each), as well as hypothetical proteins (3) and signal transduction mechanisms (2).

Interestingly, 8 genes encoding glycolytic pathway enzymes were essential in all three species. One gene encoding 2,3-bisphosphoglycerate-dependent phosphoglycerate mutase was classified as non-conclusive in *S. pyogenes* MGAS5005 [18]. The essential genes include *eno* and *gapA*, encoding enolase and glyceraldehyde 3-phosphate dehydrogenase, respectively. These enzymes not only have metabolic roles but also act as moonlighting proteins that bind host plasminogen and contribute to virulence in other streptococci [39,40]. The subcellular localisation of these enzymes in *S. anginosus* 980/01 will be investigated further.

Although the bulk of genes essential under optimal conditions are common for the three strains, we identified 40 essential genes unique to *S. anginosus* 980/01. The majority of these genes encode transcription–related proteins and hypothetical proteins with unknown function (**Fig. 4**). However, this group also includes:

- *sodA*, encoding superoxide dismutase, a key enzyme in the oxygen defence system,
- *ssaC* encoding the substrate-binding component of a manganese ABC transporter (SsaABC), essential for manganese acquisition, oxidative stress resistance, and full virulence in streptococci [41,42],
- *purB*, encoding adenylosuccinate lyase, catalyzing key reactions in the *de novo* purine biosynthesis pathway, is indispensable for nucleotide production and cellular proliferation,
- *clpC* and *clpX*, encoding ATPase subunits of the Clp protease complex, as well as *mecA*, encoding an adaptor enabling ClpC activity, and *cts*R, a regulator of the *clp* genes; all genes shown to be involved in adaptive response in *B. subtilis* [43,44].
- *ldh*, encoding L-lactate dehydrogenase, which catalyzes the conversion of pyruvate to lactate while regenerating NAD⁺, a critical step for maintaining glycolytic flux under anaerobic or microaerophilic conditions.

**Figure 4.**
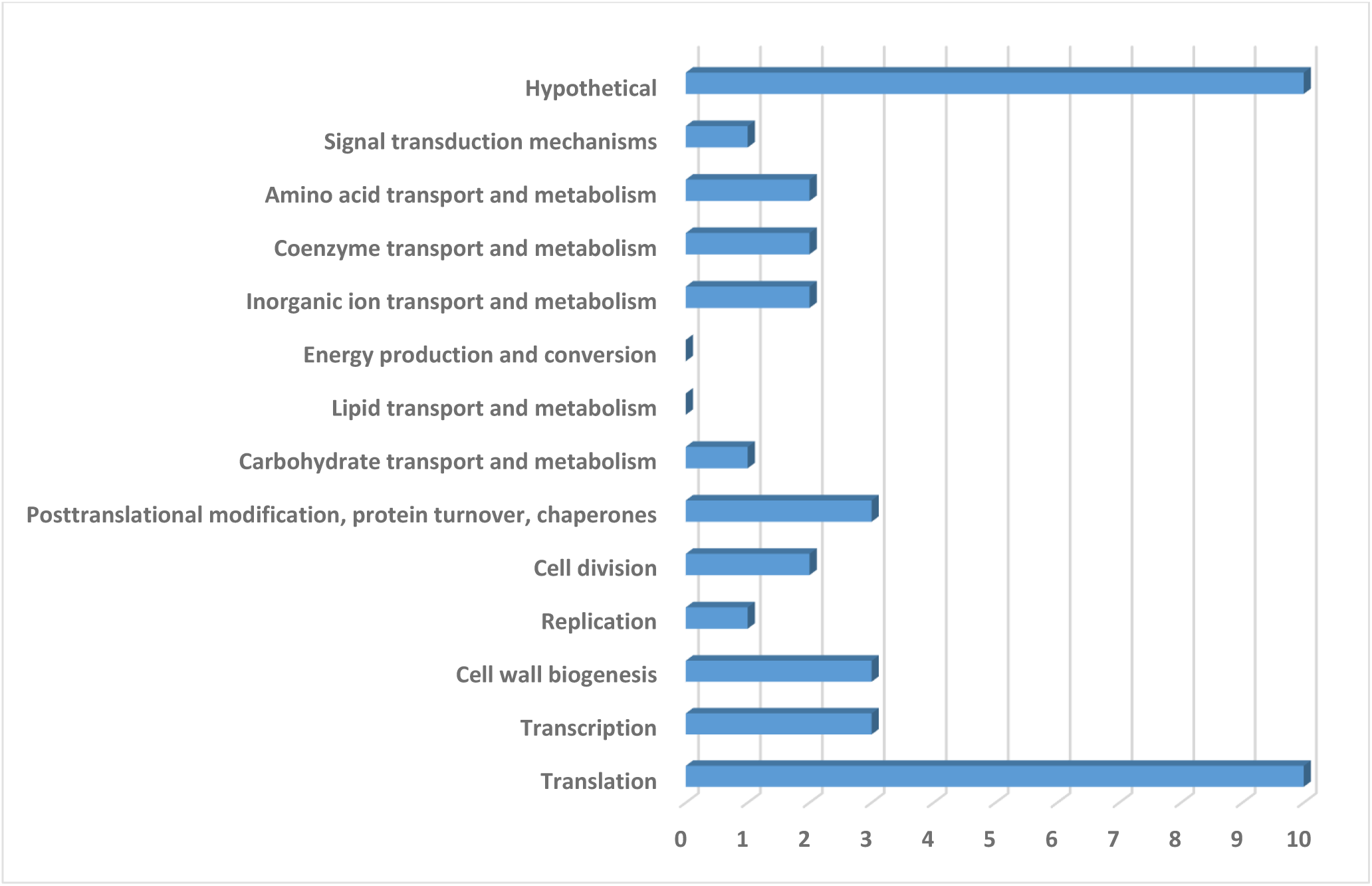
The functional classification of 40 genes uniquely essential for *S. anginosus* 980/01.

## Discussion

This study provides the genome-wide map of essential genes in *S. anginosus* 980/01, an invasive strain isolated from sepsis, with the use of high–density transposon mutagenesis (TraDIS) approach. Due to considerable genetic and phenotypic heterogeneity observed among *S. anginosus* strains [8,10], there is a clear need for molecular characterization to better understand both conserved and strain-specific requirements of this opportunistic pathogen. Despite growing recognition of the clinical relevance of the *S. anginosus* and other members of the *S. anginosus* group, functional genomic data for this species remain limited. Our findings contribute to filling this gap.

We identified 348 genes essential for growth under laboratory nutrient-rich, CO_2_-enriched aerobic conditions. Many of these genes are conserved across *S. pyogenes* MGAS5005 [18] and *S. agalactiae* A909 [29], reflecting core metabolic and cellular processes essential to streptococcal viability. However, 40 genes were uniquely essential in *S. anginosus* 980/01, highlighting species-specific physiological traits and potential therapeutic targets.

Among these, *sodA,* encoding a sole superoxide dismutase in *S. anginosus*, was identified. This correlates with the need for a CO₂-enriched atmosphere to support colony formation, suggesting limited oxidative stress tolerance. A similar dependence on *sodA* has been observed in *S. thermophilus*, where its disruption leads to oxygen hypersensitivity which can be rescued by manganese ions [45].

Also uniquely essential was *ssaC*, encoding the substrate-binding component of the SsaACB, the manganese transporter. Manganese is a key cofactor for enzymes involved in oxidative stress defense, including superoxide dismutase, however, it plays a role beyond catalytic functions (for review [46]). The essentiality of *ssaC* under nutrient-rich conditions highlights low manganese bioavailability, necessitating high-affinity uptake systems [45].

Genes involved in protein quality control, *clpC* and *clpX* encoding ATPases of the Clp protease complex, and together with *mecA* and a putative *ctsR* gene were also found to be essential in *S. anginosus* 980/01, and dispensable in *S. pyogenes* MGAS5005 and *S. agalactiae* A909. In *S. pneumoniae* R6, *clpX*, but not *clpC*, is also essential [47]. In many bacteria, impairment of the Clp protease has pleiotropic effects on cell wall composition or virulence (for review [48]). Notably, in *B. subtilis*, all four genes are involved in the heat shock response [43,44]. Given that the IS*S1-*transposon mutant library in this study was generated at 40°C, a temperature that may trigger proteotoxic stress, the observed essentiality could reflect a condition-dependent requirement for these genes.

We also found *purB*, encoding adenylosuccinate lyase, to be essential. This indicate a dependence on de novo purine synthesis, which may reflect adaptation to purine-limited environments such as mucosal surfaces or abscesses [49]. Notably, *S. agalactiae* A909 encodes two *purB* homologs, potentially providing functional redundancy and explaining its non-essentiality in that species. Given the limited data on purine availability in such niches, *purB* may represent a metabolic vulnerability worth further investigation.

Finally, *ldh*, encoding L-lactate dehydrogenase, was also uniquely essential in *S. anginosus* 980/01, reflecting its reliance on homolactic fermentation for NAD⁺ regeneration. In contrast, *S. pyogenes* and *S. agalactiae*, can employ alternative fermentation pathway as mixed-acids fermentation [38].

In summary, this study identifies essential genes in *S. anginosus* 980/01, providing insight into its metabolic constraints and potential vulnerabilities. These findings advance our understanding of *S. anginosus* physiology and provide the basis for developing targeted therapeutic strategies against this emerging pathogen. Further studies including comparisons across multiple bloodstream isolates could help pinpoint conserved targets for drug development.

## Acknowledgements

The study was funded by the National Science Centre, Poland (grant number **2018/29/B/NZ6/00624**). We extend our gratitude to Prof. Izabela Sitkiewicz (SGGW, Warsaw, Poland) for sharing the nucleotide sequence ahead of its official release.

## Data Availability Statement

The *S. anginosus* 980/01 genome sequence is available under GenBank accession number: CP183189. The datasets presented in this study have been deposited in the BioProject database under accession number PRJNA1228600.

