## Supplemental File 2 for "Genome-Wide Identification of Essential Genes in the Invasive *Streptococcus anginosus* Strain"

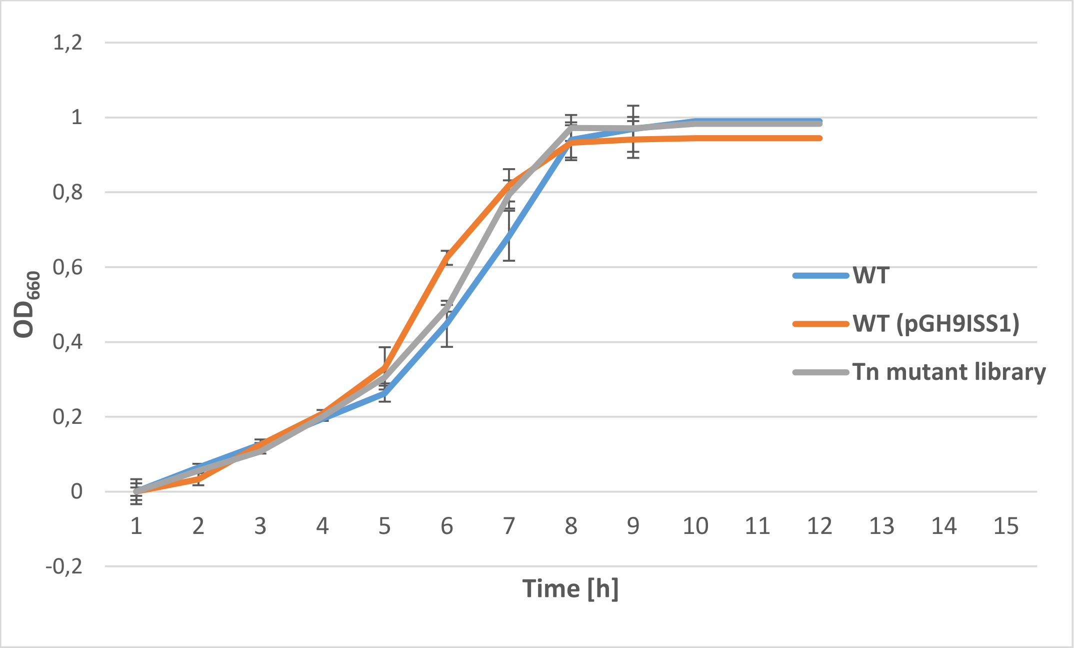


**Supplemental File S2** Growth curves of *S. anginosus* 980/01 (WT), *S. anginosus* 980/01 with pGh9:ISS*1* and ISS1 transposon mutant library of *S. anginosus* 980/01. The experiment was performed in quadruplicate, the standard deviation is marked.
